# Acidification of Cytoplasmic pH in *Escherichia coli* Provides a Strategy to Cope with Stress and Facilitates Development of Antibiotic Resistance

**DOI:** 10.1101/2020.02.24.963876

**Authors:** Esmeralda Z. Reyes-Fernández, Shimon Schuldiner

## Abstract

Awareness of the problem of antimicrobial resistance (AMR) has escalated, and drug-resistant infections are named among the most urgent issues facing clinicians today. Bacteria can acquire resistance to antibiotics by a variety of mechanisms that, at times, involve changes in their metabolic status, thus altering diverse biochemical reactions, many of them pH-dependent. In this work, we found that modulation of the cytoplasmic pH (pH_i_) of *Escherichia coli* provides a thus far unexplored strategy to support resistance. We show here that the acidification of the cytoplasmic pH is a previously unrecognized consequence of the activation of the *marRAB* operon. The acidification itself contributes to the full implementation of the resistance phenotype. We measured the pH_i_ of two resistant strains, developed in our laboratory, that carry mutations in *marR* that activate the *marRAB* operon. The pH_i_ of both strains is lower than that of the wild type strain. Inactivation of the *marRAB* response in both strains weakens resistance, and pH_i_ increases back to wild type levels. Likewise, we showed that exposure of wild type cells to weak acids that caused acidification of the cytoplasm induced a resistant phenotype, independent of the *marRAB* response. We speculate that the decrease of the cytoplasmic pH brought about by activation of the *marRAB* response provides a signaling mechanism that modifies metabolic pathways and serves to cope with stress and to lower metabolic costs.

**SIGNIFICANCE:** The decreasing effectiveness of antibiotics in treating common infections has quickened in recent years, and resistance has spread worldwide. There is an urgent need to understand the mechanisms underlying acquisition and maintenance of resistance, and here we identify a novel element in the chain of events leading to a full-fledged clinically relevant state. The *Escherichia coli* multiple antibiotic resistance (*mar*) regulon is induced by a variety of signals and modulates the activity of dozens of target genes involved in resistance to antibiotics. We report here a thus far unidentified result of this activation: acidification of the cytoplasmic pH. Manipulation of the cytoplasmic pH with weak acids and bases, independently of the *mar* response, shows that the acidification significantly increases resistance.

## INTRODUCTION

Antimicrobial resistance (AMR) is a growing public health threat of broad concern, and drug-resistant infections are named among the most urgent problems facing clinicians today (1-4). Bacteria can acquire resistance to antibiotics by horizontal gene transfer, activation of regulatory loci and genetic mutations that modify the drug target, increase drug efflux, and activate the expression of drug inactivating enzymes (5-8). High-level, clinically relevant resistant strains usually result from the accumulation of several mutations that, at times, involve changes in their metabolic status, thus altering diverse biochemical reactions (9-11). Metabolic rearrangements and changes in rates of reactions necessitate the existence of mechanisms that maintain tight homeostasis of all the physicochemical parameters in the cell within a narrow range of operation, consistent with the survival of the organism. This dynamic state of equilibrium includes many variables such as pH, and solutes and ion concentrations.

Because of the accumulating evidence on the metabolic adaptation of antibiotic-resistant cells, we focused our attention on the cytoplasmic pH of these cells. Protons play a crucial role in biochemical networks because changes in intracellular pH affect cell functioning at different levels as it impacts protein folding, enzyme activities, and the protonation of biological macromolecules, lipids, and other metabolites. Moreover, the proton electrochemical gradient across the cell membrane is key to the generation and conversion of cellular energy. It is, therefore, not surprising that the cytoplasmic pH of all living organisms is tightly controlled.

A dynamic, finely tuned balance between proton-extruding and proton-importing processes underlies pH homeostasis (12-15). Protons are extruded continuously and actively from the cytosol across the plasma membrane. The energy for proton extrusion can be provided, among others, by redox, light, or ATP driven reactions. Depending on the energy source and the environmental conditions, metabolism may produce a variety of acids, and the ensemble of membrane-bound transporters, among them cation proton exchangers, act in a coordinated mode to finely tune the cytoplasmic pH.

Numerous proteins that have crucial functions in cell physiology have been reported to be highly sensitive to small changes in the surrounding pH, and this is suggestive of a role of pH in signaling (16). A diverse collection of enzymes and transporters are transcriptionally regulated to favor acid consumption and base production at low pH (15). Moreover, changes in pH_i_ have been shown to correlate with cellular proliferation and spore germination (17, 18), to initiate and direct cell migration (19) and taxis and to trigger apoptosis. Not surprisingly, disturbances in cytoplasmic and organellar pH homeostasis, arising from either metabolic or genetic perturbations, are associated with the progression of distinct pathophysiologic states (16).

In *Escherichia coli*, a wide variety of techniques have been used to measure cytoplasmic pH (13-15). We used here a pH-sensitive derivative of the green fluorescent protein (GFP), known as ratiometric pHluorin, to measure the cytoplasmic pH of *E. coli* (20-22). EV18 and EVC are highly resistant strains that originated from *E. coli* BW25113 cells subjected to an *in vitro* lab evolution process. They carry multiple mutations that accumulated sequentially to confer high-level resistance, two to three orders of magnitude higher than the wild type strains (23, 24). Per previous findings, the wild-type cells maintain their cytoplasmic pH within a range of pH 7.4–7.7 over an external pH range of 6.0–8.1 (13-15). We find here that, under the same conditions, the resistant strain displays a substantially more acidic pH_i_ between 7.1-7.5.

We identified here one pathway involved in the reported acidification. Knockout of *marR* prevents activation of the *mar* response, restored the pH_i_ of both strains to wild type values, and significantly decreased their resistance to norfloxacin and chloramphenicol, respectively. Moreover, in parallel to the well-documented role of the *mar* regulon (8, 25), we found that activation in wild-type cells by external agents results in acidification of the cytoplasm.

Further experiments where we manipulated pH_i_ artificially supported the existence of a correlation between acidic pH_i_ and antibiotic resistance. Thus, acidification of pH_i_ in wild type cells, otherwise sensitive to norfloxacin, significantly increased their resistance. Alkalization of pH_i_ of resistant cells sensitized them to norfloxacin. The mechanism that brings about acidification and the connection between pH_i_ and resistance are two fundamental issues that deserve further analyses. Although we are only beginning to uncover the importance of pH homeostasis in antibiotic resistance, the findings described here suggest that this new player deserves further attention.

## RESULTS

### 1. Cytoplasmic pH of EV18, a strain highly resistant to norfloxacin, is lower compared to the wild-type BW25113 strain

Several studies have shown that the metabolism of antibiotic-resistant strains is altered due to adaptive changes (9-11). Since cytoplasmic pH (pH_i_) is such a central parameter for many biological processes, we explored the possibility that the metabolic alterations could also be associated with modifications of the intracellular pH of antibiotic-resistant strains. We, therefore, compared the pH_i_ of wild type and highly-resistant strains developed in our laboratory (24). To do so, we adapted a method previously described by Martinez and collaborators (20) in which a pH-sensitive derivative of the green fluorescent protein (GFP), known as ratiometric pHluorin, was used to determine the intracellular pH of *E. coli* cells (**Fig. 1A**). In our adaptation of the method, the desired strains were transformed with the pGFPR01 plasmid (harboring *pHluorin*). For the construction of a standard calibration curve, we incubated the WT strain at different pHs (5.92-8.20) in Minimal Medium A (MMA) containing 40 mM benzoic acid and 40 mM methylamine hydrochloride (collapsing compounds), a weak acid and weak base, respectively, that at high concentrations disrupt the transmembrane ΔpH resulting in the equalization of the external pH with the intracellular pH. Therefore, the indicated pH values below the graph correspond to the actual pH_i_ for each condition. The results in **Fig. 1B**, demonstrate the clear difference between the spectra exhibited at different pHs. As previously shown, the ratio (410/470 nm) monotonically increases with an increase in pH. By applying a Boltzmann sigmoid best-fit curve, we were able to use these ratios for calculating the pH_i_ for our experiments **(Fig.1C)**.

**Figure 1.**
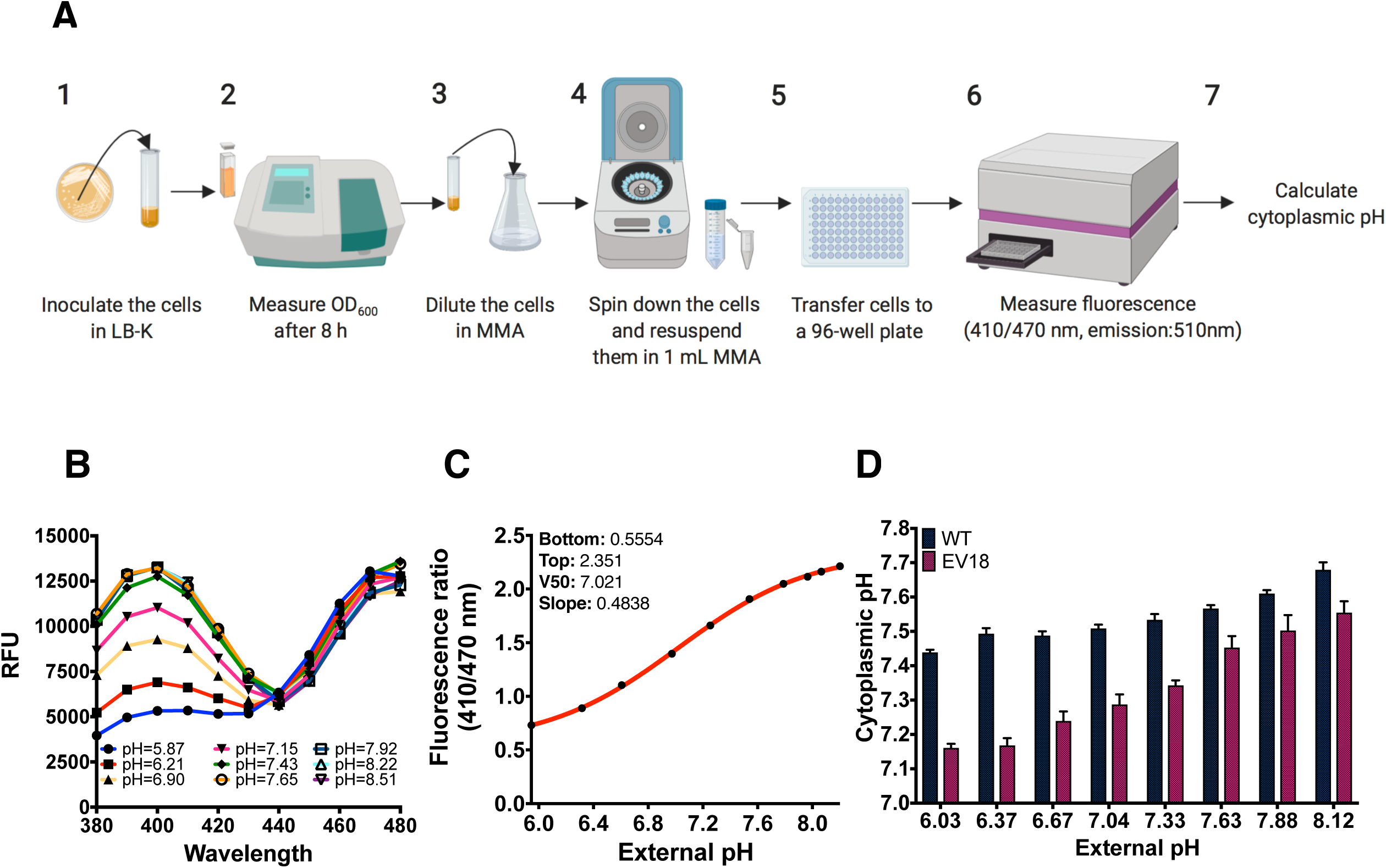
Measurement of the cytoplasmic pH (pH_i_) of the WT and EV18 strains. **(A)** Development of the method for determination of the intracellular pH in *E. coli* by using the ratiometric pH sensor pHluorin. (1) *E. coli* cells transformed with the pGFPR01 plasmid grown in LB-K medium for 8 h (2 and 3) were diluted in MMA + glycerol as a carbon source and 0.2% of arabinose and incubated ON at 25°C. (4) After spinning down and resuspension in 1 mL of MMA they were (5) transferred to a 96-well microtiter plate (OD_600_=0.1). (6) After 1 h with constant shaking (37°C) fluorescence was assessed (excitation at 410 and 470 nm with an emission at 510 nm). (7) pH_i_ values were obtained by calculating the fluorescence ratio 410/470 nm and interpolating this value in the calibration curve. **(B)** Fluorescence spectra of the WT strain (380-480 nm, emission: 510 nm) generated after incubating the strain for 1 h in MMA + glycerol at different pHs (5.92-8.20) in the presence of 40 mM benzoic acid and 40 mM methylamine hydrochloride (collapsing compounds). **(C)** Calibration curve for the measurement of pHi of the BW25113 *E. coli* strain. WT+pGFPR01 strain was incubated in a plate reader for 1 h (37°C, constant shaking) in the presence of MMA +glycerol prepared at different pHs and collapsing compounds. A Boltzmann sigmoid best-fit curve was applied to the extracted ratio data (410/470 nm) by using the GraphPad Prism 7 software. **(D)** Comparison of the cytoplasmic pH of the WT+pGFPR01 and EV18+pGFR01 strains incubated at different external pHs. Strains were incubated for 1 h (37°C, constant shaking) in MMA + glycerol at the indicated pH.

In **Fig. 1D** we see that, as expected from previous findings, *E. coli* cells tightly regulate their pH_i_ between 7.44 and 7.68 under the pH range tested (13-15). In contrast, the homeostatic pH_i_ of the EV18 strain, under the same conditions, is significantly lower, between 7.16 and 7.55. A similar behavior is displayed by the EVC strain **(Fig. S1)**. These data indicate that under the pH range tested, the pH_i_ of the resistant strains EV18 and EVC is lower than that of the WT.

### 2. Activation of the *mar* operon modifies the homeostatic value of the cytoplasmic pH

The *mar* locus confers drug resistance by altering the expression of multiple genes located in the bacterial chromosome. The *E. coli mar* locus comprises two transcriptional units: *marC* and *marRAB.* MarA, regulates the expression of a large number of chromosomal genes (the mar regulon), including those specifying MDR efflux transporters, other proteins (e.g., porins) that mediate antibiotic susceptibility and a large number of different targets, resulting in significant changes in the metabolome (10, 25-27). To test whether the acidification of pH_i_ is related to the metabolic changes produced by its induction, we inactivated the regulon by a chromosomal deletion of *marR*, the first gene in the transcriptional unit.

As expected, the deletion of the *marR* gene in both strains decreased their resistance to NF (**Fig. 2A**) and chloramphenicol **(Fig. S1B)**, respectively. EV18 was capable of growing on LB-KPi plates up to 300 μM NF to the highest dilution, while visible growth of the EV18 Δ*marR* was only possible on plates containing 100 μM NF (**Fig. 2A**). While EVC grows at concentrations as high as 200 μM, its *marR* knockout can hardly cope with 25 μM **(Fig. S1B).**

**Figure 2.**
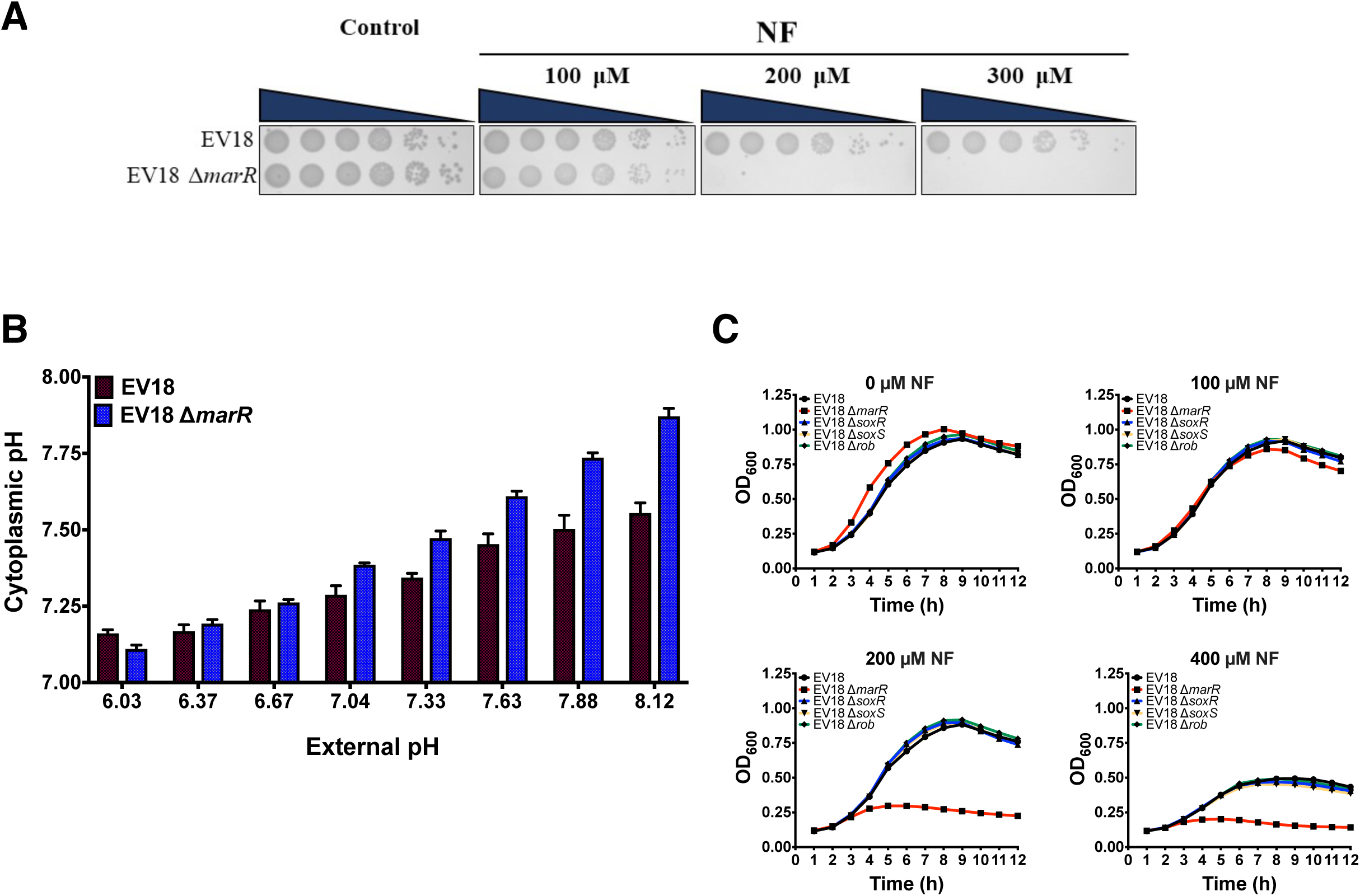
Deletion of *marR* decreases the resistance to NF and is associated to an increase in pH_i_. **(A)** Serial 10-fold dilutions of cell suspensions of the EV18 and EV18 Δ*marR* grown on LB-KPi agar supplemented with the indicated concentrations of NF. **(B)** Comparison of the cytoplasmic pH of the EV18 and EV18 Δ*marR* cells incubated at the indicated pHs. **(C)** Growth of the single mutants of *marR, soxR, soxS* and *rob* in the EV18 strain during exposure to increasing concentrations of NF for 12 h in in LB-KPi medium.

Strikingly, in parallel with the increased susceptibility to NF, we show in **Fig. 2B** and **S1A** that the pH_i_ of EV18 and EVC Δ*marR* is significantly higher than that of the parent strains. These findings imply that inactivation of the *mar* regulon diminishes the antibiotic resistance and restores the cytoplasmic pH to the wild type homeostatic value suggesting a possible correlation between both phenomena.

Since the *mar* response shares many target genes with the *sox* and *rob* regulons, we also tested whether inactivation of *soxR, soxS*, and *rob* affects sensitivity to NF. All three knockouts in EV18 display the same resistance as their parent strain showing that, under these conditions, only the *mar* response plays a significant role **(Fig. 2C)**.

In *E. coli*, expression of the *mar* operon is induced by many antimicrobial compounds but also by the seemingly unrelated salicylic acid, menadione, and others. To assess whether activation of *mar* in otherwise antibiotic sensitive cells affects pH_i_, we treated wild type BW25113 cells with menadione, an activator that does not have a direct effect on pH_i_. We measured pH_i_, resistance to NF, and levels of relevant transcripts. The results in **Fig. 3A** show that, as expected, activation of *mar* upon exposure to menadione increases resistance to NF in a dose-dependent manner. In parallel, the pH_i_ of cells treated with menadione decreases from 7.51 ± 0.01 to 7.22 ± 0.03 at a concentration of 0.63 mM and even lower to 7.03 ± 0.01 at 1.25 mM, a concentration that impairs growth (**Fig 3C**). Transcript levels of *marA* and *soxS* increase upon exposure to menadione (**Fig 3E**). Menadione is an oxidizing agent, and, as such, it also activates the *sox* operon (28, 29). To test whether its effect is due solely to the activation of *mar*, we also tested the results of exposure of a Δ*marR* strain (**Fig. 3B, D, and F**). Transcript levels of *soxS* rose to higher levels than in the wild type strain (**Fig. 3F**), and the effect on resistance and pH_i_ were similar, albeit somewhat weaker than in the wild type strain. Both regulons share a large part of their targets, and the results suggest that acidification of cytoplasmic pH and the resulting increase in resistance to NF can be attributed to the activity of targets common to both regulons.

**Figure 3.**
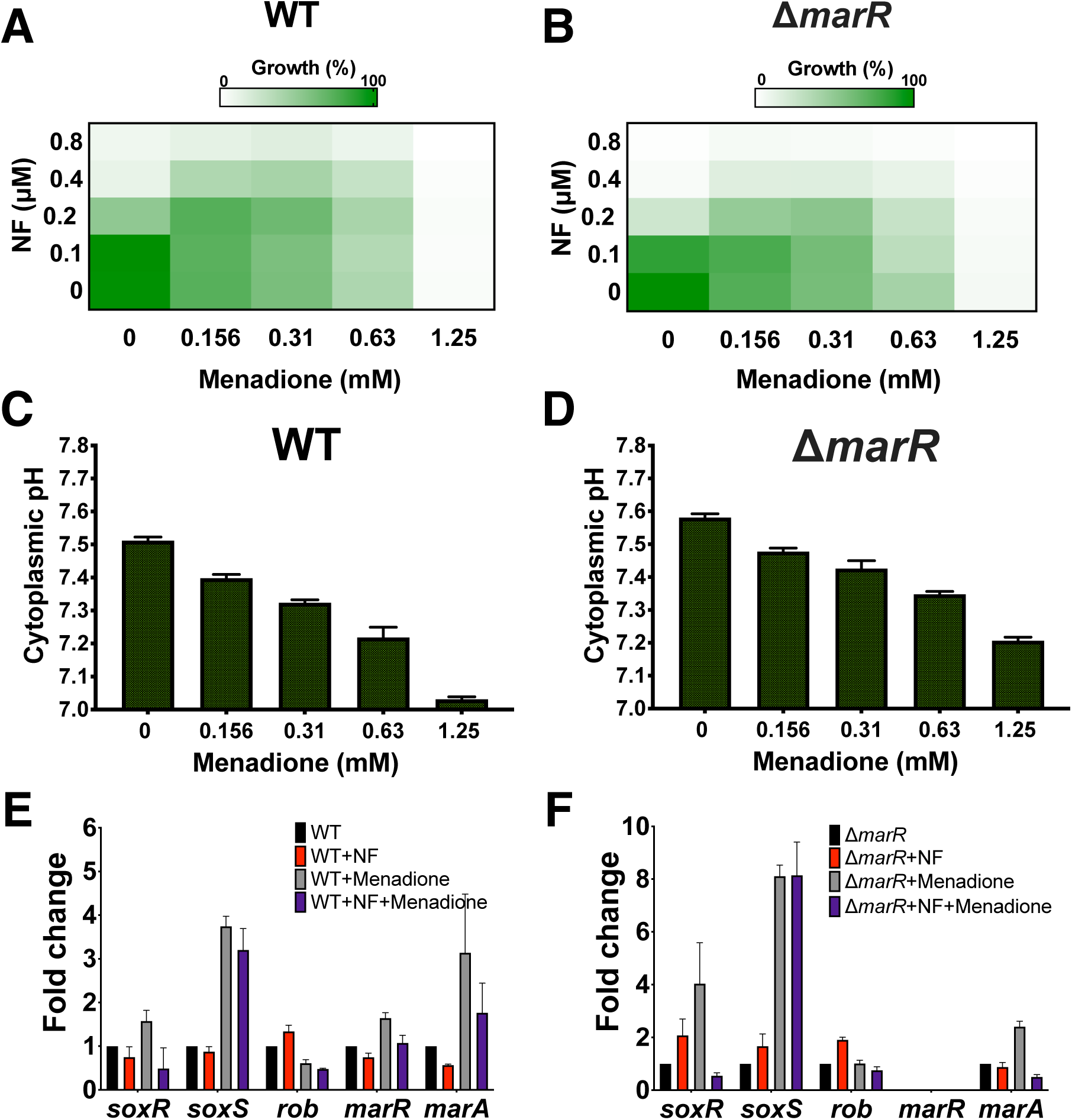
Activation of the *marR* response results in acidification of the cytoplasm. **(A-B)** Microdilution checkerboard analysis of the WT and Δ*marR* strains grown in LB-KPi medium with NF (0.1-0.8 μM) in combination with menadione. The magnitude of growth is shown as a heat plot. **(C-D)** Cytoplasmic pH of the WT and Δ*marR* strains after 1 h incubation in MMA + glycerol (pH=7.4) in the presence of increasing concentrations of menadione (0.156-1.25 mM). **(E-F)** Expression analysis of *soxR*, soxS, *rob, marR* and *marA* in the WT and Δ*marR* strains after 30 min of exposure to NF and menadione alone and in combination. mRNA expression was normalized to the gene glyceraldehyde 3-phosphate dehydrogenase (*gapDH*).

The findings presented above suggest that acidification of pH under stress conditions may be part of a general strategy that results from metabolic changes and/or regulatory modifications in the activity of membrane-bound transporters, among them cation proton exchangers, that act in a coordinated mode to finely tune the cytoplasmic pH.

### 3. Manipulation of pH_i_ with weak acids

The existence of a correlation between cytoplasmic pH and the *mar* response raises the question whether the acidification itself plays a role in resistance. To approach the subject, we manipulated the cytoplasmic pH with the aid of weak acids. They acidify the cytoplasm due to the higher permeability of the protonated species (30, 31). The latter equilibrates across the plasma membrane and releases a proton in the cytoplasm regenerating the less permeant negatively charged species (**Fig. 4B**). With the aid of weak acids, we lowered the cytoplasmic pH of wild type cells, an otherwise sensitive strain. We asked whether this manipulation provides an advantage to cope with inhibitory concentrations of antibiotics. Some of the classical acids used, salicylic (SA) and benzoic acids, are also known for their ability to induce the *marRAB* operon and antibiotic resistance. Therefore, we also analyzed the deletion mutant *marR*, incapable of activating the *marRAB* operon and, additionally, we also used sorbic acid (SoA), an aliphatic acid not known to activate the latter.

**Figure 4.**
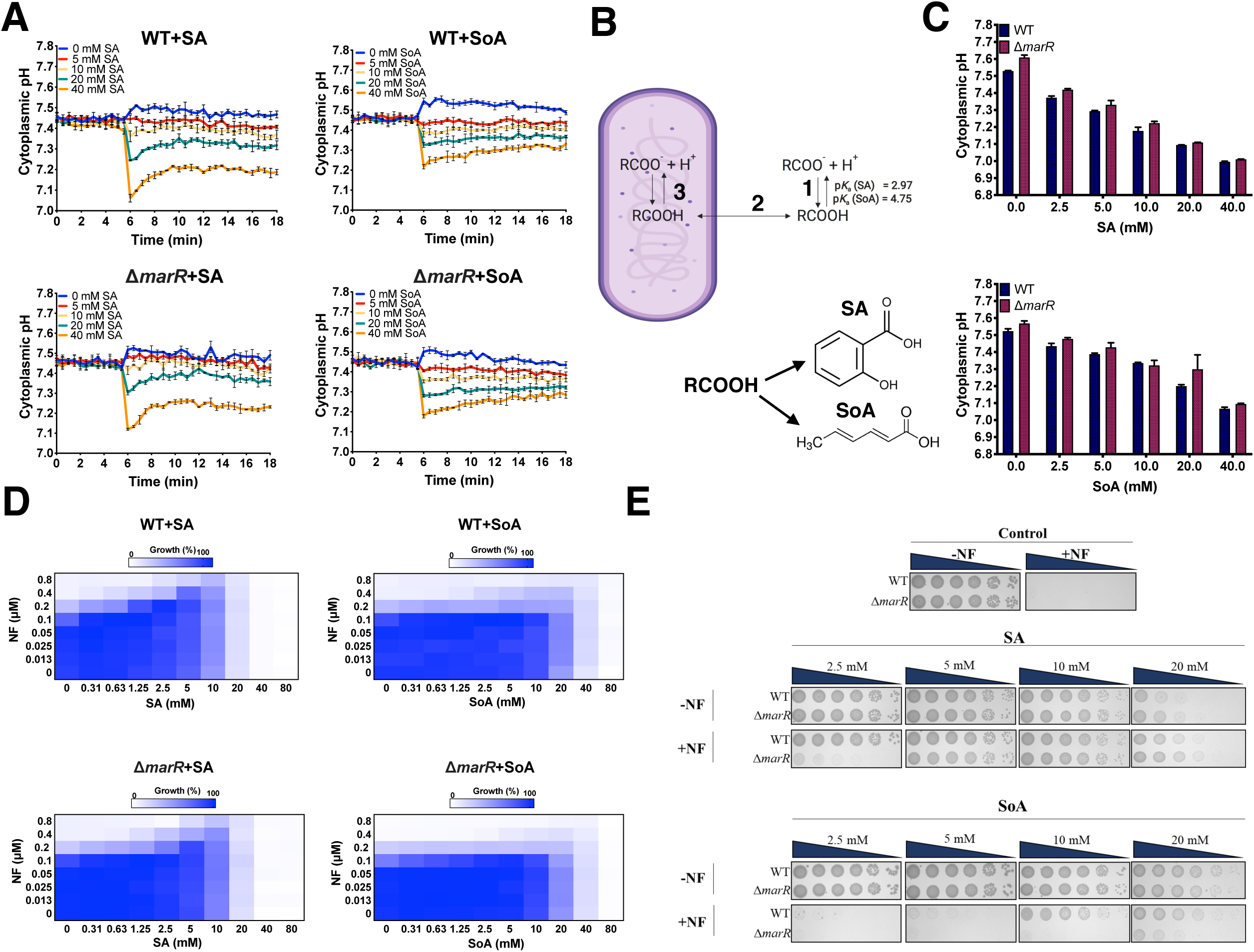
Acidification of the pH_i_ provides and advantage to cope with inhibitory concentrations of NF. **(A)** Response of the intracellular pH of the WT and Δ*marR* strains upon exposure to SA or SoA. Cells were grown in MMA +glycerol, washed and resuspended in PBS to OD_600_=0.1 (see Materials and Methods). After 6 minutes of fluorescence measurements, SA or SoA were added to the indicated concentrations. Measurements continued for an additional 12 minutes and the cytoplasmic pH was calculated with a standard calibration curve. **(B)** Acidification of the pH_i_ by weak acids. (1) The equilibrium between the protonated and unprotonated forms is determined by the individual p*K*_*a*_ of the acid. (2) The protonated and uncharged weak acid (RCOOH) diffuses faster than the charged species and equilibrates across the membrane. (3) Since the cytoplasmic pH is higher than the p*K*_*a*_ of the weak acid, this results in its dissociation (RCOO^-^ + H^+^) and acidification of the cytoplasm. The concomitant decrease in the concentration of the protonated species allows its additional permeation. **(C)** Cytoplasmic pH of the WT and Δ*marR* strains incubated in the presence of weak acids. Cells were grown in MMA with glycerol (0.5%) and pH_i_ was measured after 1 h incubation with either SA or SoA. **(D)** Microdilution checkerboard analysis of the WT and Δ*marR* strains grown in LB-KPi medium with NF in combination with either SA or SoA. The magnitude of growth is shown as a heat plot. **(E)** Serial 10-fold dilutions of cells suspensions of the WT and Δ*marR* strains grown on LB-KPi agar plates supplemented with NF and either SA or SoA. NF was added to the plates to a concentration of 0.2 μM while the effect of SA and SoA was assayed at 2.5, 5, 10 or 20 mM. Pictures were taken after 16 h incubation at 37°C.

As illustrated in **Fig. 4A** addition of SA and SoA induces a dose-dependent rapid decrease in the pH_i_ for both strains from their initial value of 7.45. The response of WT and Δ*marR* to SA and SoA was similar, but, in both cases, SA had a much potent effect than SoA. After a more prolonged incubation with weak acids in the growth medium, pH_i_ decreased from ∼7.5 to around 7.24 and 7.06 with 20 and 40 mM SA, respectively (**Fig. 4C**). The SoA mediated acidification of pH_i_ was less pronounced and reached values of 7.30 and 7.12 at 20 and 40 mM. In parallel to their effect on pH_i_, the growth rate was strongly inhibited by 20 mM SA and somewhat less by 40 mM SoA (**Fig. S2A**). In **Fig. S2B**, we show the dependence of the rate of growth on pH_i_. Growth seems to be relatively well tolerated even at pH_i_ of 7.2 (with SoA) and 7.3 (with SA), while below this value, inhibition is significant and practically full when pH_i_ drops to 7.0-7.1.

The transmembrane pH gradient is part of the proton electrochemical gradient (PMF) across the cell membrane, essential for the generation and conversion of cellular energy. Previously, it has been shown that a decrease in pH_i_ could be compensated by an increase in the membrane potential (32, 33). To rule out the possibility that the effect of the weak acids is due to a possible impact on PMF, we measured equilibrium levels of uptake of lactose in BW25113 cells (Δ*lacZ4787)* transformed with a plasmid coding for LacY. LacY is a PMF driven MFS transporter with a stoichiometry of 1H^+^/lactose (32, 34). The equilibrium levels of radiolabeled lactose provide an accurate estimate of PMF. We show the results of such an experiment in **Fig. S2C**. A slight but significant decrease of the proton motive force (PMF) (mV) from 172.51 ± 0.13 to 161.66 ± 0.51 was detected at the highest concentrations of SA but not of SoA. We conclude that at a wide range of concentrations of both acids, an increase in the transmembrane potential maintains the PMF at a stable value and supports the conclusion that the acid-mediated effects described below are due to their impact on pH_i_.

### 4. Cytoplasmic pH and antibiotic resistance in wild type

Once we showed the ability of both SA and SoA to acidify pH_i_, we tested their effect on the resistance to NF of WT and Δ*marR*. In the absence of weak acids, WT and Δ*marR* strains showed progressing growth inhibition above 0.1 μM NF and no visible growth above 0.4 μM NF **(Fig. 4D)**. Noteworthy, the addition of SA or SoA to the strains under these conditions dramatically improved their growth with full restoration at 0.2 μM NF by the addition of 2.5 mM SA and a significant improvement even as high as 0.8 μM in the presence of 5 and 10 mM SA **(Fig. 4D)**. Similarly, the incubation of WT and Δ*marR* with SoA also exhibited a noticeable increase in growth at higher concentrations of NF, even though its effect was not as substantial as the one induced by SA **(Fig. 4D).** The weaker effect of SoA is most likely due to its lower potency in acidification. The above observations were also confirmed by measuring CFU on LB-KPi agar plates with 0.2 μM NF and 2.5-20 mM of either SA or SoA **(Fig. 4E)**. As shown on the control plates (top), both strains grew alike; however, neither one was capable of growth in the presence of 0.2 μM NF. The addition of SA (2.5-10 mM) or SoA (10 mM) significantly alleviated the growth inhibition. For the Δ*marR* cells, growth with NF was fully restored at 5 and 10 mM SA, and only partially with 20 mM SoA.

We conclude that cytoplasm acidification by itself provides resistance, independent of the activation of the *marRAB* regulon.

### 5. Acidification of the cytoplasmic pH of the EV18 and EVC Δ*marR* knockouts restores most of the resistance of the parent strains

We next tested whether similar manipulations of pH_i_ in the EV18 Δ*marR* with weak acids restores some of the lost resistance of the EV18 strain. The results in **Fig. 5A** and **B** show that exposure of EV18 Δ*marR* to weak acids (±NF) caused the decline of the cytoplasmic pH, and, as shown for the wild type, also here SA was more potent than SoA **(Fig. 5A and B)**. We then tested the effect of SA and SoA on NF resistance. The strains were grown on agar plates supplemented with weak acids at the indicated concentrations in the presence and absence of NF **(Fig. 5C)**. Antibiotic concentration was 400 μM and 200 μM for the EV18 and EV18 Δ*marR*, respectively. Growth of EV18 was only partially inhibited at 400 μM, and SA and SoA only slightly improved the growth deficiency. On the other hand, EV18 Δ*marR* showed no visible growth at 200 μM NF, but increasing concentrations of SA and SoA significantly restored it. For both strains, growth was severely inhibited at 20 mM of SA without NF **(Fig. 5C)**. In a similar experiment, we show that SA also restores the resistance of EVC Δ*marR* to NF **(Fig. S1D)**. EVC is highly resistant to chloramphenicol but developed only a small concurrent resistance to NF and can grow only at around 1.6 μM NF **(Fig. S1C and D)**, and SA has only a minor effect on its ability to grow at higher concentrations. On the other hand, EVC Δ*marR* displays a higher pH_i_ and does not grow above 0.1 μM NF. SA induce acidification of the cytoplasm dramatically restores resistance to the level of EVC, its parent strain **(Fig. S1C and D)**. Interestingly, on the other hand, cytoplasm acidification has no effect on the resistance of EVC Δ*marR* to chloramphenicol **(Fig. S1C and D)**.

**Figure 5.**
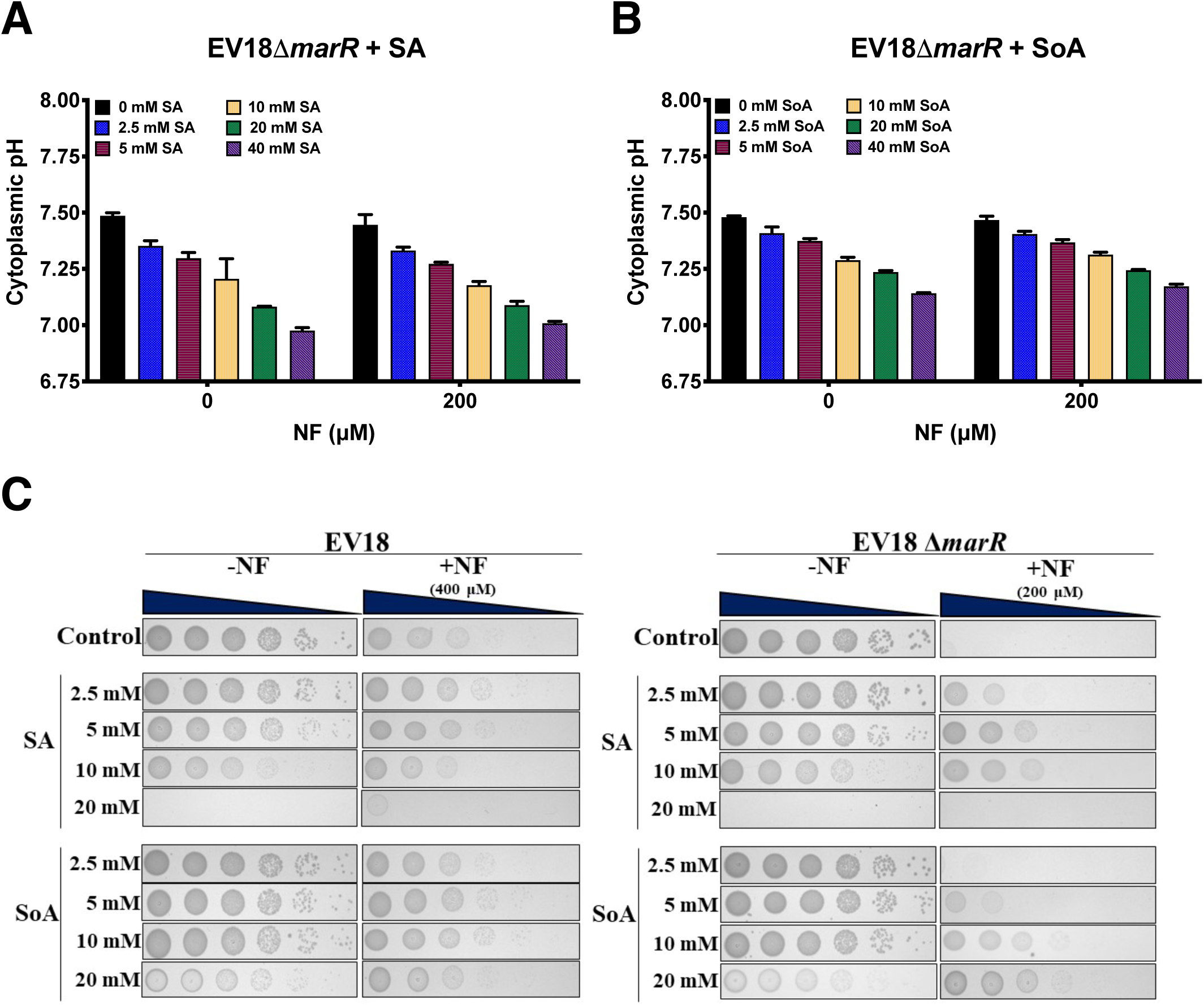
Inhibition of growth of EV18 Δ*marR* by NF is reversed by acidification of the cytoplasmic pH. **(A-B)** Determination of the cytoplasmic pH of the EV18 Δ*marR* strain incubated with NF and SA or SoA at the indicated concentrations for 1 h in MMA (37°C). **(C)** Phenotypes of the EV18 and EV18 Δ*marR* grown on LB-KPi agar plates in the presence of weak acids and NF. Agar plates were prepared with the indicated concentrations of SA or SoA. In the case of the EV18 strain (left), 400 μM of NF was added to the plates, while 200 μM of NF was used for testing the EV18 Δ*marR* strain (right). Pictures were taken after 16 h incubation at 37°C.

EV18 and EVC are highly resistant strains developed in our laboratory and have collected a considerable number of mutations that support resistance to NF and chloramphenicol, respectively (23, 24). The results thus far show that regardless of the antibiotic exposure during evolution (NF or chloramphenicol), the resistant strains have an acidic pH_i_, most likely due to the activation of the *mar* response. Thus, acidification is part of a general response to stress rather than a specific agent and is necessary for full-fledged expression of the high-level resistance. Once acidification develops, its contribution is different for antibiotics with different modes of action, in our case quinolones and chloramphenicol.

### 6. The effect of acidification is independent of the *marA/soxS/rob* regulon

The *marA/soxS/rob* regulons are involved in antibiotic resistance, superoxide resistance, and tolerance to organic solvents and heavy metals. The observations described thus far provide substantial support for the notion that the effect of acidification on resistance is independent of the activation of the *marRAB* operon. However, since many of the target genes are common to MarA, SoxS, and Rob, to test their possible involvement, we also measured the levels of transcripts of *soxR, soxS*, and *rob* **(Fig. S3).**

We measured the transcript levels of auto activator *marA*, the second gene in the operon, and detected changes of more than 400-fold when the WT strain was incubated with SA +/- NF, respectively, while, as expected, only a weaker increase of ∼ 4 (SA) and 7 (SA+NF) fold, respectively, were observed in *marA* transcripts of Δ*marR* **(Fig. S3A).** As predicted, SoA had a much weaker effect on the expression of *marA*, with an increase of ∼21 and 14-fold in the WT strain exposed to SoA and SoA+NF, respectively. No change in *marA* expression was observed under these conditions in Δ*marR* **(Fig. S3C).** In the wild type strain, *soxR* transcripts increase upon exposure to NF (∼12 fold), but there is only a minimal increase under the other conditions, and this also holds for *soxS* and *rob* transcript levels (**Fig. S3B**). In the Δ*marR* strain, there is no change in transcript levels upon exposure to SoA (**Fig. S3C**); however, SA and the combination SA and NF activate expression of *soxR* and *soxS*, most likely due to the generation of oxidative stress.

Because of the overlap of the responses, we needed further evidence to test the contention that acidification plays a direct role in resistance independent of the *marA/soxS/rob* regulon. Thus, we generated chromosomal single and double knockouts in the wild type and in EV18 cells and tested the effects of salicylic acid on growth in the presence of NF. The results in **Fig. 6A** provide, in our view, an unequivocal answer: none of the 8 mutants generated in wild type background (Δ*marR*, Δ*soxR*, Δ*soxS*, Δ*rob*, and double mutants Δ*marR*Δ*soxR*, Δ*marR*Δ*soxS*, Δ*marR*Δ*rob)* grows on plates supplemented with 0.2 μM NF. Strikingly, the addition of salicylic acid (5-10 mM) fully restores growth in all the mutants. The same knockouts were generated in the EV18 strain and challenged with 200 μM NF (**Fig. 6B**). As shown above, EV18 Δ*marR* did not grow at this concentration and, consequently, neither did the double mutants. Interestingly, EV18 Δ*soxR*, Δ*soxS, and* Δ*rob* knockouts did not affect growth, suggesting that they have no significant role in this particular strain under these conditions. Again, as shown for the wild type cells, and already described above for EV18 Δ*marR*, the addition of 5mM salicylic acid restored growth of EV18 Δ*marR* and the three double mutants.

**Figure 6.**
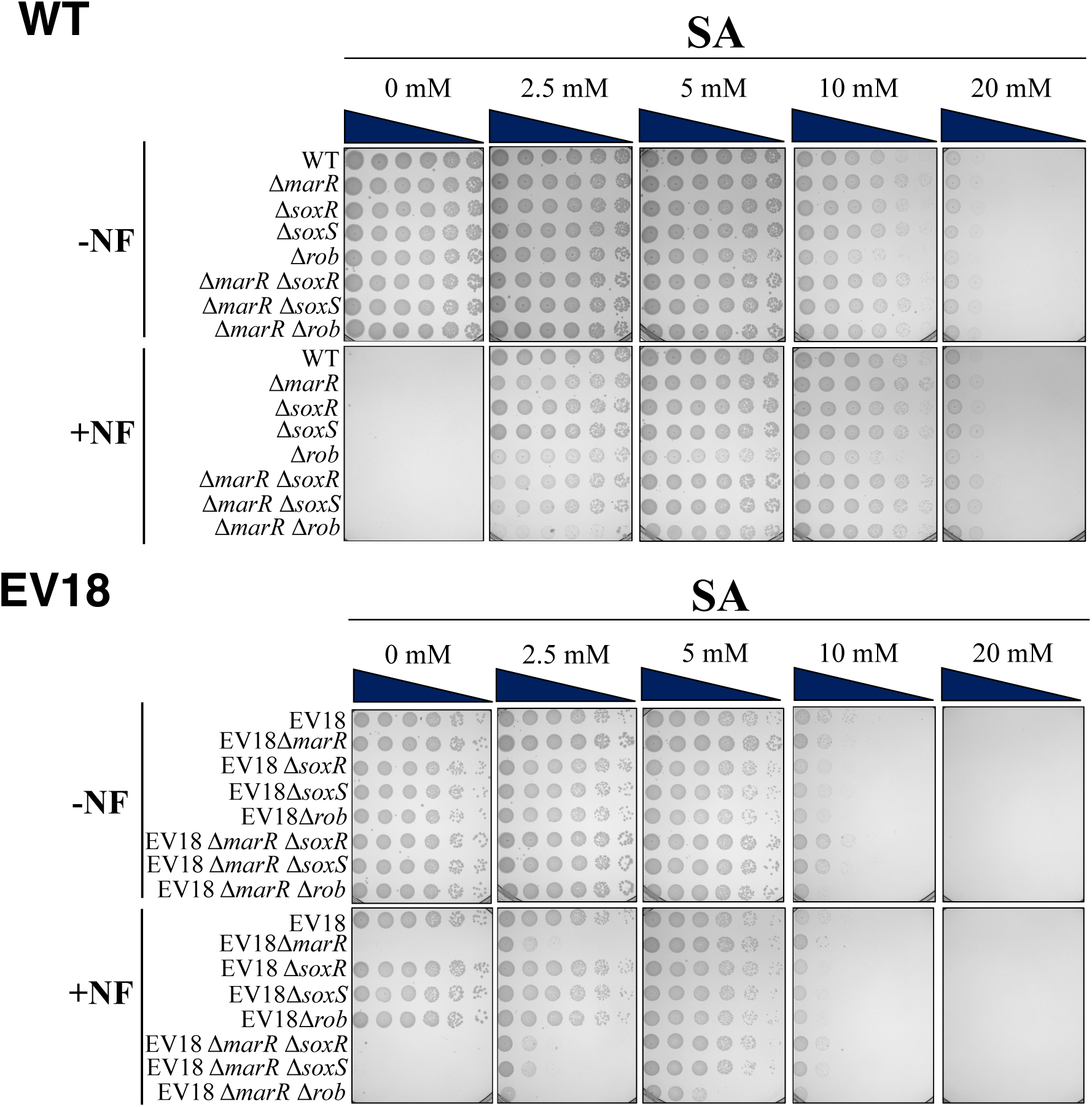
Reversal of the growth inhibition of the WT and EV18 cells exposed to high concentrations of NF is independent of *marRAB, soxRS* and *rob* functions. Serial-10 fold dilution of the single and double mutants of *marR, soxR, soxS* and *rob* in the WT **(A)** or the resistant strain EV18 **(B)** were spotted onto LB-KPi agar plates in the presence or absence of NF and increasing concentrations of SA. In the case of the WT and mutants, the plates were prepared with 0.2 μM of NF while for the EV18 strains with 200 μM of NF. Pictures were taken after 16 h incubation at 37°C.

These data provide clear support for the contention that the effect of cytoplasmic acidification on antibiotic resistance is independent of the activation of the *marRAB* as well as the *sox* and *rob* regulons. The acidification of the cytoplasm is a previously unidentified component of the response of the organism to exposure to stress such as NF, chloramphenicol, and other agents. Further work is needed to establish the mechanism that confers this resistance.

### 7. Alkalization of the pH_i_ of the resistant strains EV18 and EV18 Δ*marR* renders them more sensitive to antibiotics

Since acidification of the cytoplasm modifies the antibiotic resistance threshold so that cells are capable of coping with higher concentrations of antibiotics we asked whether the opposite effect, alkalization, could sensitize strains that are already resistant due to the accumulation of a number of mutations.

Cytoplasmic pH can be increased by the use of moderate concentrations of permeant weak bases that are not known to be metabolized. Chloroquine (CQ) increases the cytoplasmic pH of both EV18 and EV18 Δ*marR* cells in a dose-dependent manner **(Fig.7A and 7B)**. Chloroquine is a hydrophobic aminoquinoline used for the prevention and therapy of malaria with a side chain nitrogen that displays a pK_A_ of 10.1. It alkalinizes the cytoplasm due to the higher permeability of the non-protonated species **(Fig. 7C).** The latter equilibrates across the plasma membrane and binds a proton in the cytoplasm regenerating the less permeant positively charged species. At the same concentration range, in the absence of NF, CQ has a moderate effect on growth rates **(Fig. 7D and E)**. In the presence of NF, the inhibitory effect is potentiated with practically full inhibition of growth of EV18 by 0.78 mM CQ and 200 μM NF. Growth of EV18 Δ*marR* is inhibited by the combination of 100 μM NF and 0.78 mM CQ. The effect of CQ, modest but significant and reproducible, supports the notion that increasing pH_i_ sensitizes the cell to the effect of NF even in strains that accumulated many mutations that support resistance.

**Figure 7.**
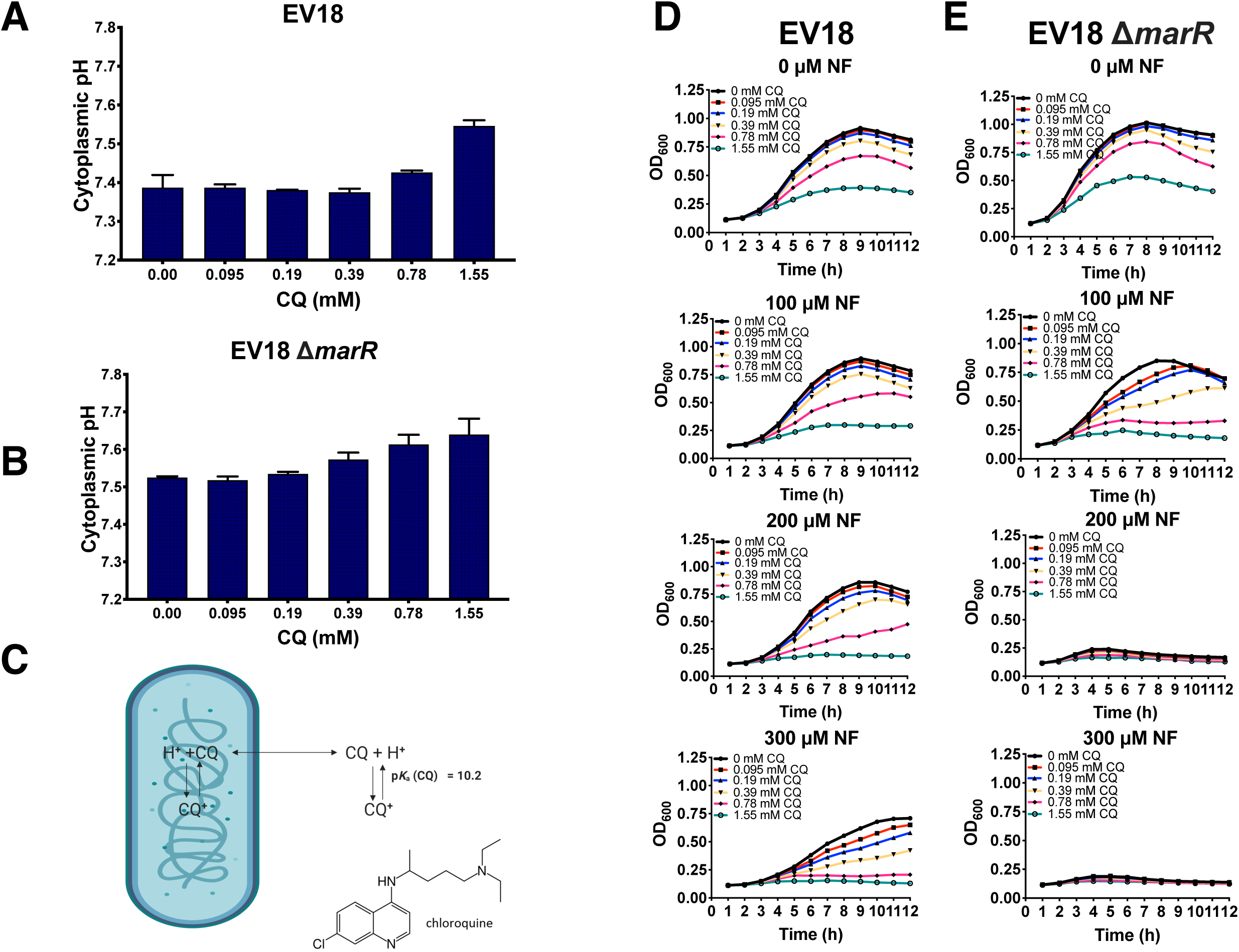
Incubation of the EV18 and EV18 Δ*marR* strains with chloroquine (CQ) increases their sensitivity to NF. **(A)** Measurement of the cytoplasmic pH of the EV18 and **(B)** EV18 Δ*marR* strains after incubation for 1 h in MMA + glycerol in the presence of increasing concentrations of CQ (0.095-1.55 mM). (**C**) Mechanism of alkalization by CQ. **(D)** Growth curves of the EV18 and **(E)** EV18 Δ*marR* generated after 12 h of incubation (37°C) with increasing concentrations of CQ and NF in LB-KPi medium.

## DISCUSSION

Intracellular pH (pH_i_) plays a crucial role in cellular physiology since it is linked to a variety of biological processes, including gene expression, energy generation, enzyme activity, and reaction rates. To the best of our knowledge, the influence of pH_i_ on antibiotic resistance has not yet been examined. Antibiotic resistance is a natural phenomenon that predates the modern selective pressure of clinical antibiotic use (35, 36). Strategies to acquire high-level clinically-relevant resistance to antibiotics are exceedingly wide-ranging, and our understanding of this phenomenon at the molecular and population levels is now enabling us to understand the details and scope of this serious health threat. Much is still to be learned, and surprises always await us. The selected mechanism of resistance by the organism depends on a multitude of parameters. Here we describe a thus far unidentified component of the response to exposure to antibiotics. We suggest that acidification of the cytoplasm provides a basal level of protection that may also act synergistically with mutations that confer resistance.

In this work, we showed that EV18 and EVC, strains highly resistant to NF and chloramphenicol, respectively, maintain a lower cytoplasmic pH compared to the sensitive wild-type strain BW25311. We presented evidence that these results may be explained by the impact in bacterial metabolism that results from activation of the *mar* response. The multiple antibiotic resistance (*mar*) locus was identified as a determinant for cross-resistance to structurally unrelated compounds by altering the expression of multiple genes (26, 37). One of the previously unrecognized consequences of this reorganization is the hereby reported acidification of the cytoplasmic pH. Thus, in this work, we showed that inactivation of the *marRAB* regulon in antibiotic-resistant cells reversed the acidification and decreased the resistance. Moreover, activation of the *mar* response by an agent such as menadione in wild type cells, otherwise sensitive to antibiotics, results in pH_i_ acidification. In a Δ*marR* strain, menadione, a known oxidizing agent, activates the *sox* response that also induces acidification of the cytoplasm. Even though each member of the *mar/sox/rob* regulon preferentially responds to a particular inducer, there is a substantial overlap in the target genes (25, 26, 37, 38). The results hint that pHi acidification may be a more general response to stress than previously imagined.

We then showed that the modulation of the intracellular pH in sensitive and resistant *E. coli* strains alters their susceptibility to antibiotics independently of the *marRAB* regulon. To differentiate between the effect of SA on *marR* and the direct acidification, we tested its effect on *marR* knockouts, and we used a weak aliphatic acid not known to interact with *marR* but still capable of acidifying the cytoplasm in a dose-dependent manner. Weak acids have been previously reported to affect the proton motive force (39, 40), but their effect is highly dependent on the external pH. Uncoupling or decrease of PMF depends on the futile leak of the charged acid, the less permeant species, and this leak is relevant only when the internal concentration is high. We performed all the work reported here at an external pH equal to the cytoplasmic pH since, under these conditions, the weak acids equilibrate between the internal and external medium but do not concentrate driven by pH gradients (13). Indeed, under these conditions, SA reduced only very slightly the proton motive force of the WT, while SoA had no impact on it.

Many studies have been performed regarding the response to low pH conditions in the environment, and it was found that *E. coli* uses a broad range of regulatory proteins to impart sophisticated control to survive exposure (41, 42). This response differs from the reaction to organic acids that act mainly by activation of the *mar* regulon and by direct effect on pHi (20, 40). Although the effect of SA on increased resistance to antibiotics has been traditionally attributed to the activation of the *mar* regulon, several studies already suggested a Mar-independent pathway. Accordingly, in the presence of salicylate, mutants in the *marRAB* operon were still able to grow at concentrations of antibiotics, otherwise non-permissive to sensitive strains (43, 44).

Changes in pH_i_ may play a direct role in the regulation of expression of some of the targets attributed to the *mar* response. These changes may also modify directly specific reactions and alter the balance between critical components such as glutathione. Acidification will moderately change the equilibrium of the latter in favor of the reduced form and provide better buffering against oxidizing agents. Methylglyoxal, a highly reactive and toxic dicarbonyl compound, is formed as a by-product of glycolysis. It was shown that, in *E. coli*, protection against methylglyoxal is mediated by cytoplasmic acidification similar in its magnitude to that reported here (45). *E. coli* possesses two glutathione-gated potassium channels, KefB and KefC, that are activated by glutathione-S-conjugates formed with methyl-glyoxal. Activation of the channels leads to cytoplasmic acidification (45, 46).

Further work is required to elucidate the mechanism by which acidification relates to increased resistance to antibiotics. The results described here provide a strong framework and novel tools for future research.

## Abbreviations

(AMR): Antimicrobial Resistance;
(SA): Salicylic acid,
(SoA): Sorbic acid,
(MMA): Minimal Medium A,
(NF): Norfloxacin

## Funding Information

This work was supported by Grant 143/16 from the Israel Science Foundation and Grants A1004 and M497-F1 from the Rosetrees Trust. The funders had no role in study design, data collection, and interpretation, or the decision to submit the work for publication

## Acknowledgment

SS is Mathilda Marks-Kennedy Emeritus Professor of Biochemistry at the Hebrew University of Jerusalem. The authors declare non-competing interests.

## MATERIALS AND METHODS

### 1. Bacterial strains, growth media, and culture conditions

Bacterial strains, plasmids, and primers used in this study are shown in **Tables S1-S4**. *Escherichia coli* strains BW25113 were grown in LB-K broth [10 g/L tryptone, 5 g/L yeast extract and 7.45 g/L KCl], LB-KPi broth [LB-K medium containing 70 mM potassium phosphate buffer, pH=7.4] or minimal medium A (MMA) (47) with 0.5% glycerol or glucose as a carbon source, and supplemented with 1X MEM Amino Acids solutions M5550 and M7145 (Sigma-Aldrich). Most of the mutants analyzed in this study were created as described by Datsenko and Wanner (48), except for Δ*soxR*, Δ*soxS*, and Δ*rob*, which were obtained from the Keio collection (49). All antibiotics and reagents in this study were purchased from Sigma-Aldrich, except for ampicillin (Amp) and sodium bicarbonate, which were acquired from Teva Pharmaceuticals (Petach Tikva, Israel) and Daejung Chemicals (Korea), respectively.

### 2. Measurement of the cytoplasmic pH in *E. coli* strains

For measuring the cytoplasmic pH, the cells bearing the pGFPR01 plasmid (kindly supplied by Prof. Joan L. Slonczewski, Kenyon College) (20) were grown for 8 h (37°C/300 rpm) in 4 mL of LB-K broth with 0.2% arabinose and Amp (100 μg/mL). For the single and double mutants of EV18 (+/-pGFR01), 50 μM of norfloxacin (NF) was added to the medium. As a control, all the corresponding strains without the plasmid were also grown in parallel under the same conditions (no Amp added). In the evening, each bacterial culture was diluted to OD_600_=0.05 in 125 mL Erlenmeyer flasks containing 25 mL of MMA pH=7.4 containing 0.5% glycerol as a carbon source and 0.2% of arabinose for induction of the plasmid. Cultures were grown overnight at 25°C (300 rpm) until they reached OD_600_=0.2-0.4. The cells were then harvested by centrifugation (4500 rpm/20 min), resuspended in 1 mL of MMA (pH=7.4), without arabinose and antibiotics, and the OD_600_ was determined. For measuring the cytoplasmic pH in all the strains, a calibration curve was constructed with cells where the ΔpH as follows. WT cells (with and without the pGFPR01 plasmid) were diluted to OD_600_=0.2 in 100 μL of MMA at different pH (5.92-8.20), transferred to a 96-well NunclonTM Delta Surface plate (Thermo Scientific) and incubated for 1 h at 37°C with constant shaking. The ΔpH of the WT cells (the difference between the external pH and internal pH) was collapsed by the addition of 100 μL of MMA at each pH containing 40 mM benzoic acid and 40 mM methylamine hydrochloride (20). Fluorescence ratio 410/470 nm (emission 510 nm) was calculated for each strain. For each fluorescence intensity obtained from the WT-pGFPR01 incubated at a certain pH, a fluorescent background was subtracted from the WT cells incubated at the same pH. A Boltzmann sigmoid best-fit curve was generated by the GraphPad Prism 7 software and used for the calculation of the cytoplasmic pH. For standard measurements, the cells of the single and double mutants of the sensitive and resistant strains were prepared as described above without the addition of the collapsing compounds. For kinetic measurements the plate was incubated for 1 h at 37°C (constant shaking) in a microplate reader. As a next step, the reader measured continuously for 6 min the fluorescence intensity at 410 and 470 nm (510 nm of emission) of each well to determine the cytoplasmic pH before treatment. When the measurements finished, the plate reader was ejected and 10 μL of the stocks solutions and 2-fold dilutions of the tested addition were added to the corresponding well. As a control, 10 μL of PBS was added to the non-treated samples. The plate was placed back to the reader to measure fluorescence for another 12 min.

Strains were analyzed in triplicates, and cytoplasmic pH was calculated using the calibration curve previously constructed. All experiments were repeated at least twice.

### 3. Assays of Bacterial susceptibility to added reagents

For analyzing the sensitivity of the WT and resistant strains in *E. coli*, their phenotypes were monitored for growth rate in liquid and CFU in solid media in the presence of increasing concentrations of antibiotics and the corresponding reagent additions. Pre-cultures of the strains to be tested were grown after overnight in 4 mL of growth in LB-K (37°C/300 rpm), and the next morning cells were diluted to 1:100 in either LB-K or MMA (pH=7.4, 0.5% glycerol). Cells continued growing for several hours (until the early logarithmic phase), and they were diluted in either LB-K or MMA to a final OD_600_=0.02 for examining their growth in liquid media. Duplicate serial dilutions at a factor of 2 were made in 96-well plates for NF with each of the reagents. As a control, two wells were used to monitor the growth without antibiotics or reagents, and one well only with medium to rule out contamination. Growth was followed overtime for 16 h, measuring the optical density at 600 nm every hour, at 37°C and constant shaking in a Synergy 2 Microplate Reader (Bio Tek). Results were obtained by comparing the percentage of growth of the tested strain grown without antibiotics/reagents with the strain grown in the presence of them. Growth curves and microdilution checkerboard analysis were obtained by using the GraphPad Prism 7 software. For the phenotypes on solid medium, the cells were grown as indicated above but instead of diluting them to an OD_600_=0.02, they were brought to OD_600_=0.2 and 5 μL of 10-fold serial dilutions of the cells were spotted on LB-KPi (pH=7.4) agar plates containing increasing concentrations of antibiotics and other reagents according to the experimental setup. Plates were then incubated at 37°C overnight, and the growth of the strains was recorded by using the Fusion Fx ECL and Gel Documentation System (VILBER). Biological replicates were included for each experiment.

### 4. qRT-PCR analysis

WT and Δ*marR* cells were inoculated in 4 mL of LB-K overnight at 37°C with constant shaking (300 rpm). The next morning the cells were diluted into 4 mL fresh LB-K to an OD_600_ of 0.05 and grown until the cells reached the OD_600_ =0.4. At this point, weak acids (10 mM SA or SoA) or menadione (0.156 mM) were added to the corresponding cultures. Similarly, NF was added either alone (0.2 μM final concentration) or in combination with weak acids or menadione. Afterward, the cells were incubated back for an additional 30 min and harvested by centrifugation (3 min, 13000 rpm). Total RNA was isolated according to the protocol accompanying the PureLink® RNA Mini Kit (Ambion) and treated with DNase I (Invitrogen) to remove trace amounts of DNA. For the synthesis of cDNA, 2 μg of RNA was used per sample in a final reaction volume of 20 μL with the High Capacity RNA-to-cDNATM kit (Applied BiosystemsTM). Real-time PCR reactions were prepared in a final volume of 20 μL using the Fast SYBR® Green Master Mix (Applied BiosystemsTM), the corresponding cDNA to each sample, and the primer combinations described on **Table S3**. Reactions were run for 40 cycles (standard curve program) in experimental duplicate using two independent cDNA preparations in the StepOne Plus Real-Time PCR system (Applied BiosystemsTM). Expression levels of each queried gene were normalized using the gene *gapDH* as an endogenous control. Analysis of the gene expression was performed according to the comparative CT method (50).

### 5. Lactose transport

Uptake of [14C] Lactose of the indicated *E. coli* strains was adapted from the protocol previously described (51) with some modifications. *E. coli* BW25113 cells (Δ*lacZ4787)* transformed with pKK223/lacY, a plasmid coding for LacY under the control of the *tac* promoter, were grown overnight in MMA with 0.5% glucose (pH=7.0) at 37°C with constant shaking. Cultures were then diluted to OD_600_=0.1 in fresh MMA and grown until they reached an OD_600_ of 0.4. Subsequently, IPTG (2mM) was added for induction, and the cultures continued growing for an additional 1.5 h. The cells were then harvested (3,220 x g, 10 min) at room temperature and resuspended in pre-warmed wash buffer [100 mM potassium phosphate (pH=7.5), 10 mM MgSO4 and 0.5% glucose]. Cells were then washed once and concentrated to a final OD_600_ of 12 in wash buffer. Afterward, 25 μL of resuspended cells were placed into a glass reaction tube 12 x75 mm and kept at 37°C. The transport reaction was initiated by the addition of 0.4 mM [14C] Lactose, specific activity of 185 x 10^6^ Bq/mmole (5 mCi/mmole). This reaction was terminated at the indicated intervals by the addition of 2 mL of cold wash buffer followed by immediately passing the cells through a 25 mm GF/C filter (Whatman) and additional 2 ml washing. Reaction tubes were supplemented with extra oxygen throughout the experiment. Radioactivity was measured by the Tri-Carb®2900TR Liquid Scintillation Analyzer (PerkinElmer®). Experiments were carried out in duplicates and repeated at least twice. Concentrations of SA and SoA used for this experiment are detailed in **Fig. S2F**.

### 6. Software

The processing of the graphs was performed with GraphPad Prism 7 software. The images describing the method for measuring the pH_i_ in *E. coli* cells, as well as the figures of the mechanisms shown in this work, were created with BioRender.com.

## Figure Legends

**Figure S1. Inactivation of the *marRAB* operon in the highly resistant strain to Cam, EVC, causes an increase in the pH**_**i**_ **and sensitivity of the strain to Cam and NF.**

**(A)** Cytoplasmic pH of EVC, and its *marR* mutant. Cells were incubated for 1 h in MMA at 37°C. **(B)** Growth curves of the EVC and EVC Δ*marR* incubated for 12 h in LB-KPi with increasing concentrations of Cam (25-400 μM). **(C)** Microdilution checkerboard analysis of the EVC or **(D)** EVC Δ*marR* incubated with weak acids in the presence of either NF or Cam. Growth was recorded after 12 hours of incubation of both strains.

**Figure S2. Effect of weak acids (SA and SoA) on the rate of growth (OD**_**600**_**/h) and proton motive force (PMF) in the WT strain. (A)** Correlation between the concentration of weak acid and the growth rate in the WT strain. Cells were grown in MMA + glycerol with the indicated concentrations of weak acids. **(B)** Cytoplasmic pH vs. growth rate of the WT strain in the presence of weak acids. Concentrations of SA (blue) and SoA (red) are indicated on each graph. **(C)** Proton motive force of the WT strain in the presence of SA or SoA. Proton motive force was estimated from equilibrium values of accumulation of [^14^ C] Lactose as described in Materials and Methods. Experiments were carried out in duplicates and repeated at least twice.

**Figure S3. Real-time PCR analysis of the gene expression of *marR, marA, soxR, soxS* and *rob* in the WT and Δ*marR* strains exposed to NF, SA and SoA. (A)** qRT-PCR analysis of *marR* and *marA* mRNA levels in the WT strain after the transient exposure of the cells to SA, SoA or NF, as well as in combination of these compounds. **(B)** Expression analysis of *soxR, soxS* and *rob* in the WT strain after exposure to NF and the weak acids alone and in combination. **(C)** mRNA transcript levels of *marR, marA, soxR, soxS* and *rob* of the Δ*marR* under the same conditions as described above for the WT strain. WT and mutant strains were grown in LB-K medium until OD_600_=0.4 and exposed for 30 min to NF and weak acids. mRNA expression was normalized to the gene glyceraldehyde 3-phosphate dehydrogenase (*gapDH*).

## REFERENCES

1. K. Bush et al., Tackling antibiotic resistance. Nature reviews 9, 894–896 (2011).

2. J. Carlet et al., Society’s failure to protect a precious resource: antibiotics. Lancet 378, 369–371 (2011).

3. Center for disease control (2013) Antibiotics resistance threats in the USA, 2013. In Annual report of the CDC, Centers for disease control, USA.

4. R. Laxminarayan et al., Antibiotic resistance-the need for global solutions. The Lancet infectious diseases 13, 1057–1098 (2013).

5. J. M. Blair, M. A. Webber, A. J. Baylay, D. O. Ogbolu, L. J. Piddock, Molecular mechanisms of antibiotic resistance. Nature reviews 13, 42–51 (2015).

6. H. Nikaido, Multidrug Resistance in Bacteria. Annu Rev Biochem 78, 119–146 (2009).

7. G. D. Wright, Molecular mechanisms of antibiotic resistance. Chem Commun (Camb) 47, 4055–4061 (2011).

8. M. N. Alekshun, S. B. Levy, Molecular Mechanisms of Antibacterial Multidrug Resistance. Cell 128, 1037–1050 (2007).

9. M. Zampieri, M. Zimmermann, M. Claassen, U. Sauer, Nontargeted Metabolomics Reveals the Multilevel Response to Antibiotic Perturbations. Cell reports 19, 1214–1228 (2017).

10. M. Zampieri et al., Metabolic constraints on the evolution of antibiotic resistance. Mol Syst Biol 13, 917 (2017).

11. D. J. Dwyer, J. J. Collins, G. C. Walker, Unraveling the physiological complexities of antibiotic lethality. Annu Rev Pharmacol Toxicol 55, 313–332 (2015).

12. T. A. Krulwich, G. Sachs, E. Padan, Molecular aspects of bacterial pH sensing and homeostasis. Nature reviews 9, 330–343 (2011).

13. E. Padan, S. Schuldiner, Intracellular pH regulation in bacterial cells. Methods Enzymol 125, 337–352 (1986).

14. E. Padan, D. Zilberstein, S. Schuldiner, pH homeostasis in bacteria. Biochim Biophys Acta 650, 151–166 (1981).

15. J. L. Slonczewski, M. Fujisawa, M. Dopson, T. A. Krulwich, Cytoplasmic pH measurement and homeostasis in bacteria and archaea. Advances in microbial physiology 55, 1-79, 317 (2009).

16. J. R. Casey, S. Grinstein, J. Orlowski, Sensors and regulators of intracellular pH. Nature Reviews Molecular Cell Biology 11, 50–61 (2010).

17. B. M. Swerdlow, B. Setlow, P. Setlow, Levels of H+ and other monovalent cations in dormant and germinating spores of Bacillus megaterium. J Bacteriol 148, 20–29 (1981).

18. J. W. A. van Beilen, S. Brul, Compartment-specific pH monitoring in Bacillus subtilis using fluorescent sensor proteins: a tool to analyze the antibacterial effect of weak organic acids. Frontiers in microbiology 4, 157–157 (2013).

19. S. P. Denker, D. L. Barber, Cell migration requires both ion translocation and cytoskeletal anchoring by the Na-H exchanger NHE1. J Cell Biol 159, 1087–1096 (2002).

20. K. A. Martinez, 2nd et al., Cytoplasmic pH response to acid stress in individual cells of Escherichia coli and Bacillus subtilis observed by fluorescence ratio imaging microscopy. Appl Environ Microbiol 78, 3706–3714 (2012).

21. M. J. Mahon, pHluorin2: an enhanced, ratiometric, pH-sensitive green florescent protein. Advances in bioscience and biotechnology (Print) 2, 132–137 (2011).

22. G. Miesenbock, D. A. De Angelis, J. E. Rothman, Visualizing secretion and synaptic transmission with pH-sensitive green fluorescent proteins. Nature 394, 192–195 (1998).

23. N. Alon Cudkowicz, S. Schuldiner, Deletion of the major Escherichia coli multidrug transporter AcrB reveals transporter plasticity and redundancy in bacterial cells. PLoS One 14, e0218828 (2019).

24. Y. Shuster, S. Steiner-Mordoch, N. Alon Cudkowicz, S. Schuldiner, A Transporter Interactome Is Essential for the Acquisition of Antimicrobial Resistance to Antibiotics. PLoS One 11, e0152917 (2016).

25. T. M. Barbosa, S. B. Levy, Differential expression of over 60 chromosomal genes in Escherichia coli by constitutive expression of MarA. J Bacteriol 182, 3467–3474 (2000).

26. M. N. Alekshun, S. B. Levy, The mar regulon: multiple resistance to antibiotics and other toxic chemicals. Trends in Microbiology 7, 410–413 (1999).

27. N. Weston, P. Sharma, V. Ricci, L. J. V. Piddock, Regulation of the AcrAB-TolC efflux pump in Enterobacteriaceae. Res Microbiol 169, 425–431 (2017).

28. V. Duval, I. M. Lister, MarA, SoxS and Rob of Escherichia coli - Global regulators of multidrug resistance, virulence and stress response. Int J Biotechnol Wellness Ind 2, 101–124 (2013).

29. R. G. Martin, J. L. Rosner, Promoter Discrimination at Class I MarA Regulon Promoters Mediated by Glutamic Acid 89 of the MarA Transcriptional Activator of *Escherichia coli*. Journal of Bacteriology 193, 506 (2011).

30. J. L. Slonczewski, R. M. Macnab, J. R. Alger, A. M. Castle, Effects of pH and repellent tactic stimuli on protein methylation levels in Escherichia coli. J Bacteriol 152, 384–399 (1982).

31. S. Ramos, S. Schuldiner, H. R. Kaback, The use of flow dialysis for determinations of deltapH and active transport. Methods Enzymol 55, 680–688 (1979).

32. D. Zilberstein, S. Schuldiner, E. Padan, Proton Electrochemical Gradient in Escherichia coli Cells and Its Relation to Active Transport of Lactose. Biochem 18, 669–673 (1979).

33. S. Ramos, S. Schuldiner, H. R. Kaback, The electrochemical gradient of protons and its relationship to active transport in Escherichia coli membrane vesicles. Proc Natl Acad Sci USA 73, 1892–1896 (1976).

34. H. R. Kaback, A chemiosmotic mechanism of symport. Proc Natl Acad Sci U S A 112, 1259–1264 (2015).

35. V. M. D’Costa et al., Antibiotic resistance is ancient. Nature 477, 457–461 (2011).

36. B. Spellberg et al., The Epidemic of Antibiotic-Resistant Infections: A Call to Action for the Medical Community from the Infectious Diseases Society of America. Clinical Infectious Diseases 46, 155–164 (2008).

37. P. Sharma et al., The multiple antibiotic resistance operon of enteric bacteria controls DNA repair and outer membrane integrity. Nature communications 8, 1444 (2017).

38. R. G. Martin, J. L. Rosner, Genomics of the marA/soxS/rob regulon of Escherichia coli: identification of directly activated promoters by application of molecular genetics and informatics to microarray data. Mol Microbiol 44, 1611–1624 (2002).

39. O. H. Setty, R. W. Hendler, R. I. Shrager, Simultaneous measurements of proton motive force, delta pH, membrane potential, and H+/O ratios in intact Escherichia coli. Biophysical Journal 43, 371–381 (1983).

40. K. E. Creamer et al., Benzoate- and Salicylate-Tolerant Strains of Escherichia coli K-12 Lose Antibiotic Resistance during Laboratory Evolution. Appl Environ Microbiol 83 (2017).

41. J. W. Foster, Escherichia coli acid resistance: tales of an amateur acidophile. Nature reviews 2, 898–907 (2004).

42. U. Kanjee, W. A. Houry, Mechanisms of acid resistance in Escherichia coli. Annu Rev Microbiol 67, 65–81 (2013).

43. S. P. Cohen, S. B. Levy, J. Foulds, J. L. Rosner, Salicylate Induction of Antibiotic Resistance in Escherichia-coli - Activation of the mar Operon and a mar-Independent Pathway. Journal of Bacteriology 175, 7856–7862 (1993).

44. T. Wang, I. El Meouche, M. J. Dunlop, Bacterial persistence induced by salicylate via reactive oxygen species. Scientific reports 7, 43839 (2017).

45. G. P. Ferguson, D. McLaggan, I. R. Booth, Potassium channel activation by glutathione-S-conjugates in Escherichia coli: protection against methylglyoxal is mediated by cytoplasmic acidification. Mol Microbiol 17, 1025–1033 (1995).

46. J. Healy et al., Understanding the structural requirements for activators of the Kef bacterial potassium efflux system. Biochemistry 53, 1982–1992 (2014).

47. B. Davies, E. Mingioli, Mutants of Escherichia coli requiring methionine or vitamin B12. J Bacteriol 60, 17–28 (1950).

48. K. A. Datsenko, B. L. Wanner, One-step inactivation of chromosomal genes in Escherichia coli K-12 using PCR products. Proc Natl Acad Sci U S A 97, 6640–6645 (2000).

49. T. Baba et al., Construction of Escherichia coli K-12 in-frame, single-gene knockout mutants: the Keio collection. Mol Syst Biol 2, 2006.0008 (2006).

50. T. D. Schmittgen, K. J. Livak, Analyzing real-time PCR data by the comparative C(T) method. Nat Protoc 3, 1101–1108 (2008).

51. S. Brill, O. S. Falk, S. Schuldiner, Transforming a drug/H+ antiporter into a polyamine importer by a single mutation. Proc Natl Acad Sci U S A 109, 16894–16899 (2012).

