## Supplementary material for "Acidification of Cytoplasmic pH in *Escherichia coli* Provides a Strategy to Cope with Stress and Facilitates Development of Antibiotic Resistance": Supplememntal figures and Tables

Fig.S1

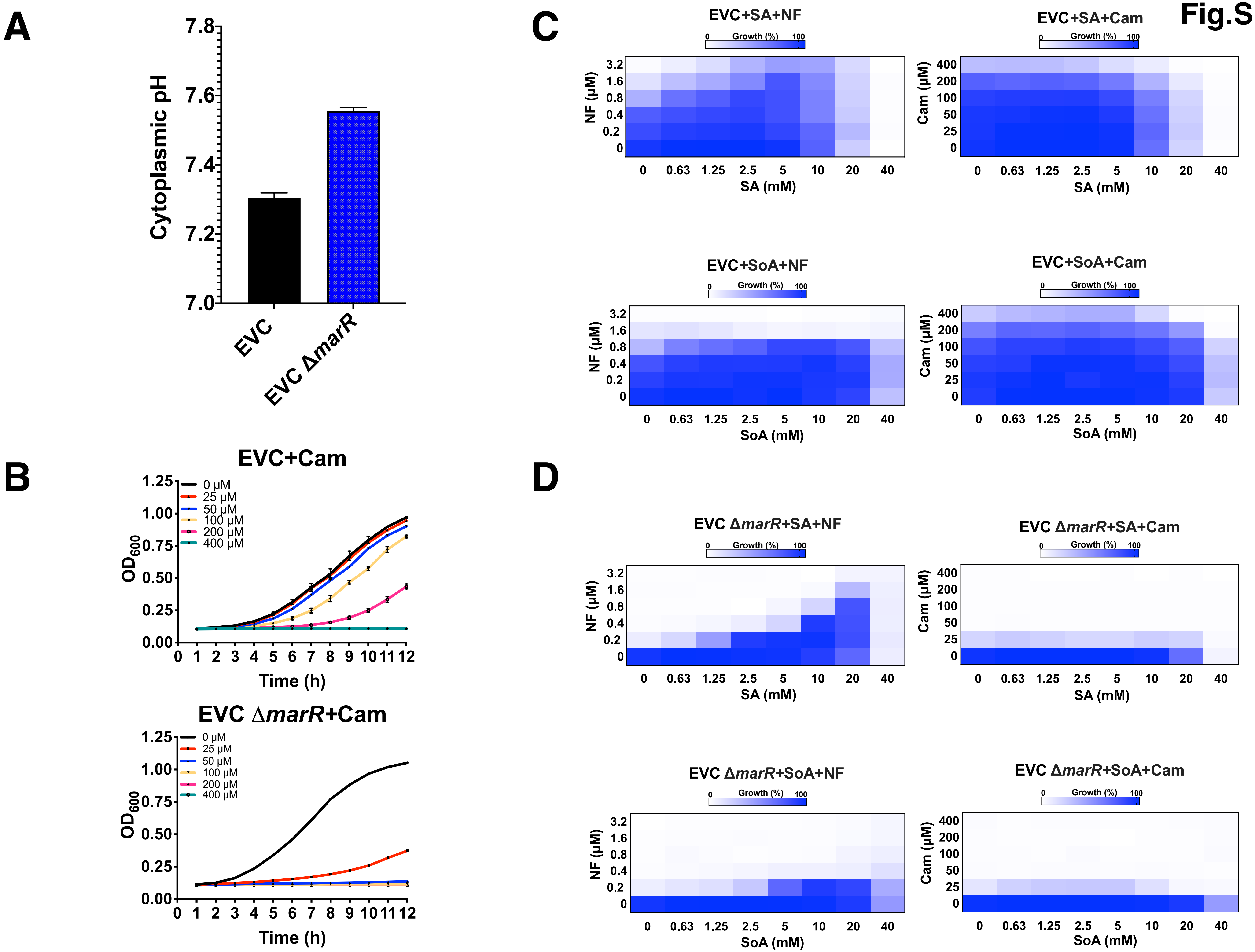
A
Cytoplasmic pH
7.8
7.6
7.4
7.2
7.0
EVC
EVC ΔmarR
B
EVC+Cam
OD600
Time (h)
0 μM
25 μM
50 μM
100 μM
200 μM
400 μM
EVC ΔmarR+Cam
C
EVC+SA+NF
EVC+SA+Cam
EVC+SoA+NF
EVC+SoA+Cam
Growth (%)
NF (μM)
Cam (μM)
SA (mM)
SoA (mM)
D
EVC ΔmarR+SA+NF
EVC ΔmarR+SA+Cam
EVC ΔmarR+SoA+NF
EVC ΔmarR+SoA+Cam

**Fig.S2**

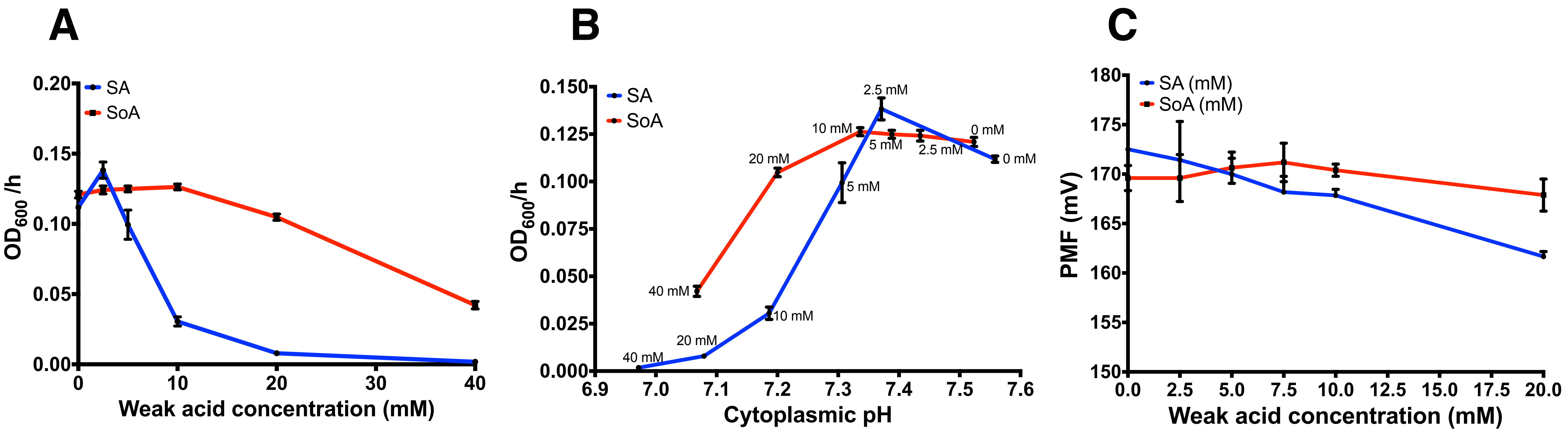
A
OD600 /h
SA
SoA
0.20
0.15
0.10
0.05
0.00
0
10
20
30
40
Weak acid concentration (mM)
B
OD600/h
SA
SoA
0.150
0.125
0.100
0.075
0.050
0.025
0.000
6.9
7.0
7.1
7.2
7.3
7.4
7.5
7.6
Cytoplasmic pH
2.5 mM
10 mM
0 mM
5 mM
2.5 mM
0 mM
20 mM
5 mM
40 mM
10 mM
20 mM
40 mM
C
PMF (mV)
SA (mM)
SoA (mM)
180
175
170
165
160
155
150
0.0
2.5
5.0
7.5
10.0
12.5
15.0
17.5
20.0
Weak acid concentration (mM)

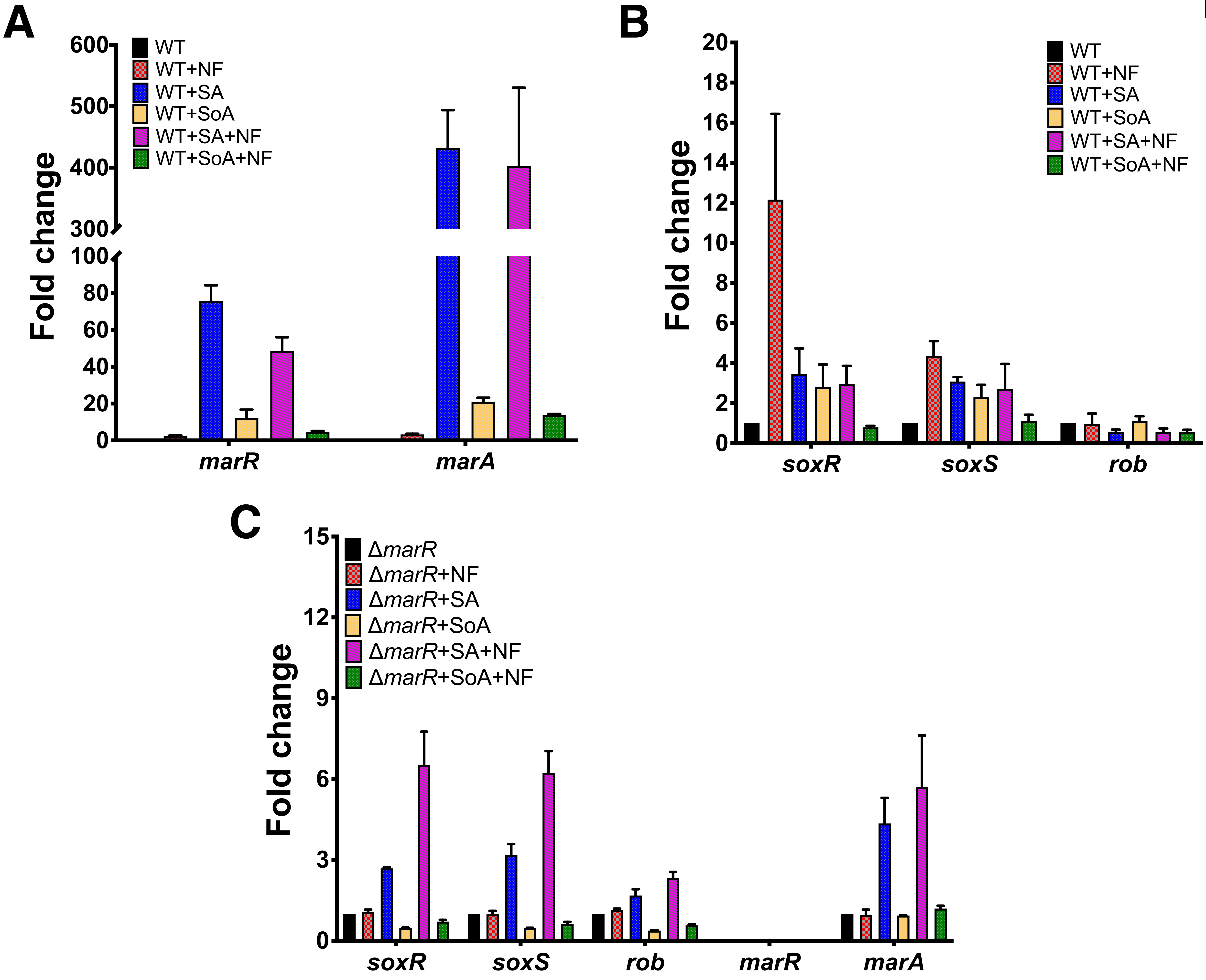
A
600
500
400
300
100
80
60
40
20
0
Fold change
WT
WT+NF
WT+SA
WT+SoA
WT+SA+NF
WT+SoA+NF
marR
marA
B
20
18
16
14
12
10
8
6
4
2
0
Fold change
WT
WT+NF
WT+SA
WT+SoA
WT+SA+NF
WT+SoA+NF
soxR
soxS
rob
C
15
12
9
6
3
0
Fold change
ΔmarR
ΔmarR+NF
ΔmarR+SA
ΔmarR+SoA
ΔmarR+SA+NF
ΔmarR+SoA+NF
soxR
soxS
rob
marR
marA

**Table S1**. ***E. coli*** **strains used for this study.**

| **Strain** | **Description** | **Reference** |
|---|---|---|
| WT | Wild-type strain (BW25113) | This study |
| Δ*marR*::*kan* | Deletion mutant of the *marR* gene in the background of the WT strain. | This study |
| Δ*soxR*::*kan* | Deletion mutant of the *soxR* gene in the background of the WT strain. | (1) |
| Δ*soxS*::*kan* | Deletion mutant of the *soxS* gene in the background of the WT strain. | (1) |
| Δ*rob*::*kan* | Deletion mutant of the *rob* gene in the background of the WT strain. | (1) |
| Δ*marR* Δ*soxR*::*kan* | Deletion mutant of the *soxR* gene in the background of the Δ*marR* strain. | This study |
| Δ*marR* Δ*soxS*::*kan* | Deletion mutant of the *soxS* gene in the background of the Δ*marR* strain. | This study |
| Δ*marR* Δ*rob*::*kan* | Deletion mutant of the *rob* gene in the background of the Δ*marR* strain. | This study |
| EV18* | NF resistant strain created by evolution *in vitro* (18 days habituation). | (2) |
| EV18Δ*marR*::*kan* | Deletion mutant of the *marR* gene in the background of the EV18 strain. | This study |
| EV18Δ*soxR*::*kan* | Deletion mutant of the *soxR* gene in the background of the EV18 strain. | This study |
| EV18Δ*soxS*::*kan* | Deletion mutant of the *soxS* gene in the background of the EV18 strain. | This study |
| EV18Δ*rob*::*kan* | Deletion mutant of the *rob* gene in the background of the EV18 strain. | This study |
| EV18Δ*marR* Δ*soxR*::*kan* | Deletion mutant of the *soxR* gene in the background of the EV18Δ*marR* strain. | This study |
| EV18Δ*marR* Δ*soxS*::*kan* | Deletion mutant of the *soxS* gene in the background of the EV18Δ*marR* strain. | This study |
| EV18Δ*marR* Δ*rob*::*kan* | Deletion mutant of the *rob* gene in the background of the EV18Δ*marR* strain. | This study |

*Resequencing of this strain revealed a previously undetected mutation in *marR.* Starting at nucleotide 329 of marR, 38 bp deletion that results in permanent activation of the *mar* regulon.

**Table S2**. **Plasmids used in this study.**

| Plasmid | Reference |
|---|---|
| pKD13 | (3) |
| pKD46 | (3) |
| pGFPR01 | (4) |
| pKKmarA | This study |

**Table S3**. **Primers used for real-time PCR in this study.**

| Gene | Primer | Sequence (5'-3') | Tm (°C) |
|---|---|---|---|
| ***gapDH*** | *gapDH*_fw_qRTPCR | ACTTACGAGCAGATCAAAGC | 52.6 |
| | *gapDH*_rv_qRTPCR | AGTTTCACGAAGTTGTCGTT | 52.4 |
| ***soxR*** | *soxR*_fw_qRTPCR | ATCCGTAACAGCGGCAATCA | 57.1 |
| | *soxR*_rv_qRTPCR | TCGCACTTAACGTATGCCCT | 56.5 |
| ***soxS*** | *soxS*_fw_qRTPCR | GACCTGGGTTATGTCTCGCA | 56.8 |
| | *soxS*_rv_qRTPCR | TTACAGGCGGTGGCGATAAT | 56.6 |
| ***rob*** | *rob*_fw_qRTPCR | CGGCGAAAGCAGGTTATTCC | 56.6 |
| | *rob*_rv_qRTPCR | TTTCGACAAACGACGAGCAC | 55.9 |
| ***marA*** | *marA*_fw_qRTPCR | CATAGCATTTTGGACTGGAT | 50.3 |
| | *marA*_rv_qRTPCR | TACTTTCCTTCAGCTTTTGC | 50.9 |
| ***marR*** | *marR*_fw_qRTPCR | CTGTAAAGGCTGGGTGGAAAG | 55.9 |
| | *marR*_rv_qRTPCR | GGTCCTGGCCAACTAATTGATG | 56 |

**Table S4**. **Primers used for generating the knock-out mutants used in this study.**

| Primer* | Sequence (5'-3') | Tm (°C) |
|---|---|---|
| ***marR*_H1P1** | GCAAAACGTGGCATCGGTCAATTCATTCATTTGACTTAT ACTTGCCTGGGATTCCGG | 71.4 |
| ***marR*_H2P2** | CGATCCAGTCCAAAATGCTATGAATGGTAATAGCGTCA GTATTGCGTCTGT GTAGGC | 70.4 |
| ***marR*_L1** | GCTAGCCTTGCATCGCATTGAA | 58.4 |
| ***marR*_L2** | GACACTTTCTCCAGTGACAGTG | 55.4 |
| ***soxR*_H1P1** | CTGTTGGGGAGTATAATTCCTCAAGTTAACTTGAGGTA AAGCGATTTATGATTCCGGGGATCCGTCGACC | 69.2 |
| ***soxR*_H2P2** | AAAACAAACTAAAGCGCCCTTGTGGCGCTttaGTTTTGT TCATCTTCCAGTGTAGGCTGGAGCTGCTTCG | 71.3 |
| ***soxR*_L1** | GTCAATATGCTCGTCAATCC | 51.1 |
| ***soxR*_L2** | CATACAATTAAAGCATCAAC | 44.8 |
| ***soxS*_H1P1** | CCCCAACAGATGAATTAACGAACTGAACACTGAAAAGA GGCAGATTTATGATTCCGGGGATCCGTCGACC | 70.1 |
| ***soxS*_H2P2** | GCGCGGGAGTTAACGCGCGGGCAATAAAATTACAGGC GGTGGCGATAATCTGTAGGCTGGAGCTGCTTCG | 73.6 |
| ***soxS*_L1** | GGTTAGCAGCGCTTTAATGC | 54.9 |
| ***soxS*_L2** | GCGTCGAAACTGAGGAGCAG | 58.4 |
| ***rob*_H1P1** | AATTACCTGATGTCAGGTGCTCGTTGTTGAAAGGATGA GGATATTTTATGATTCCGGGGATCCGTCGACC | 69.9 |
| ***rob*_H2P2** | GACGCCCCTGCATTAGATGAGCTGCAGCGTAAACGACG GATCGGAATCAGTGTAGGCTGGAGCTGCTTCG | 73.1 |
| ***rob*_L1** | CGTCAAGCCCTAAAACATAC | 51.2 |
| ***rob*_L2** | TAACTGTTCTATTTCGCGCG | 53.1 |
| **K1** | CAGTCATAGCCGAATAGCCT | 54.1 |
| **K2** | CGGTGCCCTGAATGAACTGC | 59.2 |
| **KT** | CGGCCACAGTCGATGAATCC | 58.4 |

*The primers are named as described in (3).

1. T. Baba *et al.*, Construction of Escherichia coli K-12 in-frame, single-gene knockout mutants: the Keio collection. *Mol Syst Biol* **2**, 2006.0008 (2006).
2. Y. Shuster, S. Steiner-Mordoch, N. Alon Cudkowicz, S. Schuldiner, A Transporter Interactome Is Essential for the Acquisition of Antimicrobial Resistance to Antibiotics. *PLoS One* **11**, e0152917 (2016).
3. K. A. Datsenko, B. L. Wanner, One-step inactivation of chromosomal genes in Escherichia coli K-12 using PCR products. *Proc Natl Acad Sci U S A* **97**, 6640-6645 (2000).
4. K. A. Martinez, 2nd *et al.*, Cytoplasmic pH response to acid stress in individual cells of Escherichia coli and Bacillus subtilis observed by fluorescence ratio imaging microscopy. *Appl Environ Microbiol* **78**, 3706-3714 (2012).
